# Active late Holocene accretion of mesophotic coral-rubble fields in the northern Red Sea

**DOI:** 10.64898/2026.10.07.757296

**Authors:** Gal Eyal, David K. Weinstein, Nicholas M. Hammerman, Yonatan Shaked, Adi Torfstein

## Abstract

Mesophotic coral ecosystems are commonly interpreted as thin biological veneers over relict shallow-water reef structures, but their capacity to generate late Holocene carbonate deposits remains poorly quantified. Here we combine diver excavations, rubble-bed thickness surveys, radiocarbon age-depth modelling, sediment grain-size analyses, coral-rubble assemblages, trace-element geochemistry, and modern *Alveopora allingi* growth measurements from two 40-65 m deep coral-rubble fields in the northern Gulf of Eilat/Aqaba, Red Sea. Rubble deposits are at least 1.0-1.5 m thick and accumulated at mean rates of 2.21 ± 0.52 mm yr^-1^ at the Dekel site and 1.33 ± 0.13 mm yr^-1^ at the IUI site, implying persistent carbonate accumulation over centuries to >1 kyr. Age-depth models indicate quasi-continuous deposition despite pronounced faunal changes, including a shift from *A. allingi*-dominated rubble to upper layers enriched in *Euphyllia paradivisa* at Dekel. The dominant modern rubble-forming coral, *A. allingi*, extends at 3.04 ± 0.87 mm yr^-1^ and has skeletal density of 1.17 ± 0.27 g cm^-3^. First-order cover-based accretion estimates at present coral cover fall well below the radiocarbon-derived rates, indicating that long-term mesophotic rubble accumulation integrates historical cover, secondary calcification and sediment retention rather than present live coral cover alone. These data demonstrate that mesophotic rubble fields can act as active carbonate factories and palaeoecological archives, challenging the assumption that modern mesophotic habitats are geomorphologically passive.

## 1. Introduction

Coral reefs are biogenic carbonate factories that influence shelf morphology, sediment budgets, coastal protection, and the long-term preservation of ecological and environmental change (Hubbard and Dullo, 2016; Perry and Alvarez-Filip, 2019). The accretionary record of reefs provides information on sea-level history, palaeoceanography, sedimentary processes, reef-community transitions, and anthropogenic impacts (Roff et al., 2013, 2015; Leonard et al., 2018; Yan et al., 2019). Although Holocene shallow-water reefs have been intensively investigated (Dullo, 2005; Roff et al., 2015; Hubbard and Dullo, 2016), the capacity of mesophotic coral ecosystems (MCEs), generally occurring at 30-150 m depth, to generate and preserve carbonate deposits remains poorly constrained (Loya et al., 2016; Sherman et al., 2019; Turner et al., 2019).

A persistent geological view is that many modern MCEs represent thin living veneers over older shallow-water reef foundations or antecedent hardgrounds. This interpretation is supported in some settings by ship-based cores and drowned reef records, but it is difficult to generalize because modern mesophotic habitats are rarely sampled with the spatial control needed to resolve active depositional processes (Fairbanks, 1989; Camoin et al., 2007; Woodroffe et al., 2010; Yokoyama et al., 2011; Abbey et al., 2013). Cenozoic records show that mesophotic corals can form substantial carbonate accumulations, yet depth attribution in fossil systems is often uncertain and modern analogues remain sparse (Bosellini, 1998; Morsilli et al., 2012; Novak et al., 2013; Pomar et al., 2017).

Coral rubble is central to this problem. Rubble, defined here as unattached dead coral skeletons modified by bioerosion, sediment infill, secondary calcification, cementation, and possible transport, can dominate reef deposits and act as both a taphonomic archive and a substrate for renewed coral colonization (Blanchon et al., 1997; Pandolfi and Greenstein, 1997; Rasser and Riegl, 2002; Weinstein et al., 2014, 2016; Hubbard and Dullo, 2016). Mesophotic rubble fields are widely reported, but few studies have linked their stratigraphy, age structure, sedimentology, geochemistry, and modern coral growth to quantify whether they are active depositional systems or relict sedimentary veneers.

The northern Gulf of Eilat/Aqaba (GoEA), Red Sea, provides a strong test case because mesophotic rubble fields occur as low-relief accumulations at 40-65 m depth and host living unattached corals with distinctive growth forms (Eyal et al., 2016, 2019; Eyal-Shaham et al., 2016; Weinstein et al., 2021). We use diver excavations, rubble-thickness surveys, radiocarbon age-depth models, rubble assemblages, sediment granulometry, trace-element geochemistry, and in situ growth measurements to address three questions: (1) how thick are these deposits and how fast have they accumulated; (2) do coral-rubble assemblages and sediments record ecological or environmental transitions; and (3) are modern coral growth and cover sufficient to explain the observed late Holocene accretion?

## 2. Materials and methods

### 2.1. Site setting and modern coral cover

Fieldwork was conducted in 2018-2019 at two mesophotic coral-rubble fields in the northern GoEA (Fig. 1). Site 1, offshore Dekel Beach (Dekel; 29°32′11″ N, 34°56′51″ E), occurs on a gently declining slope at approximately 45 m depth and hosts a mixed modern assemblage including unattached *Euphyllia paradivisa* and *Alveopora allingi*. The area is intermittently affected by flash floods from local wadis, which can deliver terrigenous sediment and reduce water quality (Katz et al., 2015; Eyal et al., 2016). Site 2, approximately 5 km SSW of Site 1, offshore from the Interuniversity Institute for Marine Sciences (IUI; 29°29′58″ N, 34°55′01″ E), lies on a relatively flat upper mesophotic terrace and is dominated by unattached *Alveopora* spp. (Eyal-Shaham et al., 2016; Weinstein et al., 2021).

**Figure 1.**
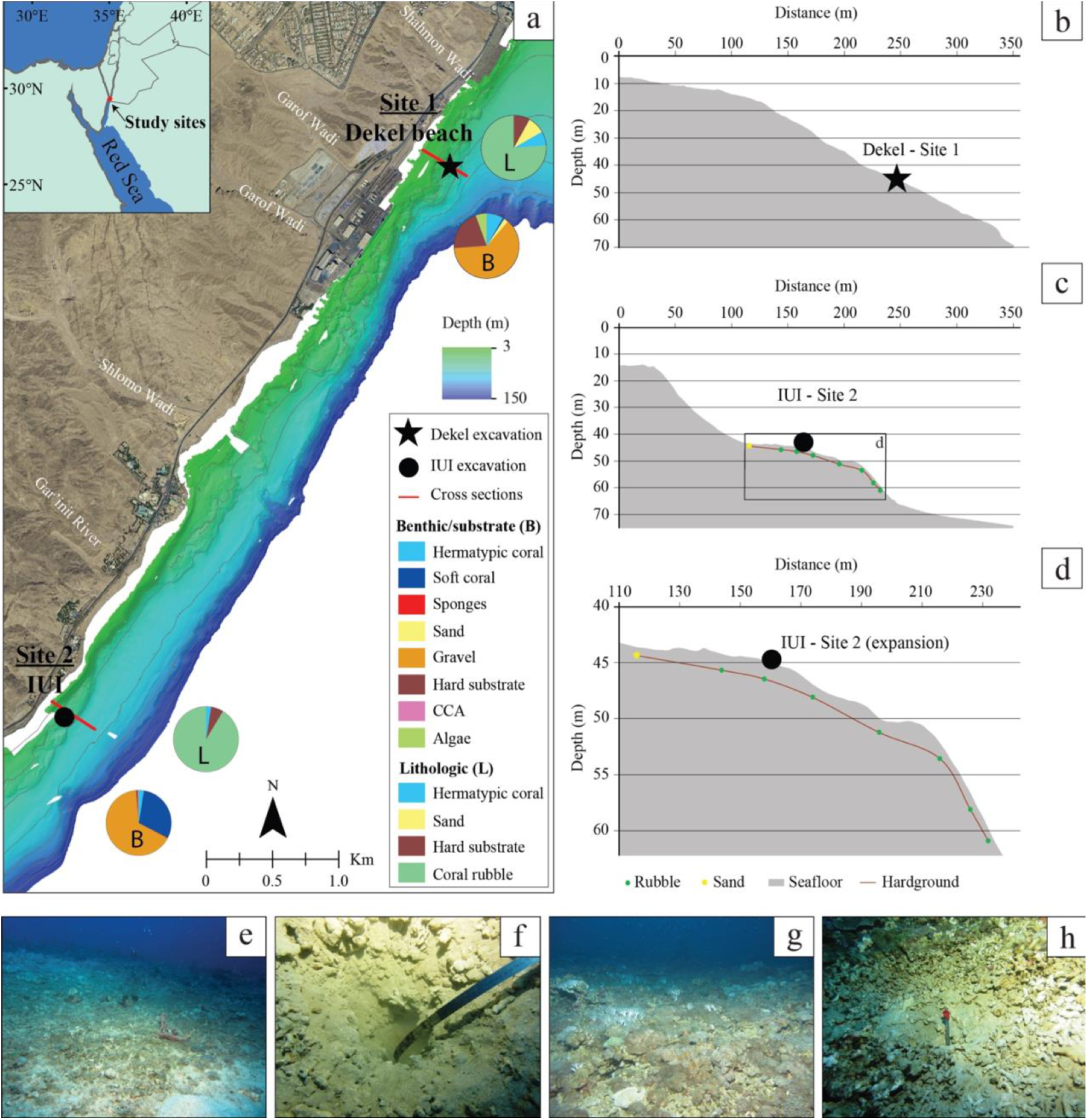
Excavation sites of two mesophotic coral-rubble fields (CRFs) in the northern Gulf of Eilat/Aqaba, Israel. (a) Map of site locations with bathymetry of shallow to mesophotic reef depths along the Israeli coast of the Red Sea. Pie charts near excavation markers represent relative contemporary benthic taxa or substrate (B) and lithologic characteristics (L) of the seabed near the excavation sites at 45 m depth. (b-c) Cross sections along the depth gradient from shallow bathymetric data to approximately 70 m depth. (d) Close-up of the cross-section from panel c highlighting the presumed subsurface hardground and rubble accumulation along the terrace. (e-f) In situ photographs of the CRF and excavation pit at Dekel Beach (Site 1). (g-h) In situ photographs of the CRF and excavation pit near IUI (Site 2).

Live coral cover and lithology were quantified from annotated photographs using random point intercept classification in CoralNet, with 100 points per image (Beijbom et al., 2015). Nine adjacent 50 × 70 cm photoquadrats were collected along a 10 m transect at Site 1 by technical divers following the 45 m contour (Tamir et al., 2019). Ten photographs near Site 2 were collected using a drop-camera system along the IUI mesophotic terrace.

### 2.2. Rubble thickness, excavations, and sample preparation

Rubble-field thickness was estimated by hammering a 2 m marked metal rebar perpendicularly into the rubble bed until refusal. Measurements were collected every 10 m along a 70 m transect across the 40-65 m depth gradient at Site 2 and near the 45 m excavation site at Site 1. At both sites, excavation pits were dug by hand and with metal shovels at 45 m depth. Rubble was recovered in approximately 10 cm depth bins, and only material confidently assigned to its original depth was retained.

Samples were sun-dried, sorted, and identified to species or genus where possible. Each taxon in each depth bin was weighed to estimate relative rubble assemblage composition. Specimens fully overgrown by secondary calcifiers were sectioned to identify internal taxa. All depth bins were inspected under a binocular microscope for bioerosion, sediment infill, taphonomic alteration, and secondary accretion. Representative pristine skeletal fragments were selected for radiocarbon dating and trace-element geochemistry. Approximately 1-5 g of skeletal material was cut, crushed to approximately 0.5 mm, cleaned following modified procedures of Clark et al. (2014), and hand-picked under a binocular microscope to isolate pristine aragonite.

### 2.3. Radiocarbon dating and age-depth modelling

Ten coral-rubble subsamples were selected for radiocarbon dating; one stratigraphically inconsistent young age was excluded because secondary aragonite needles were observed. Dating was conducted at the Poznań Radiocarbon Laboratory and Beta Analytic. Pre-bomb ages were calibrated with Marine20 using a local ΔR of 3 ± 67 yr, calculated from published known-age *Porites lutea* samples from the northern GoEA (Felis et al., 2000; Heaton et al., 2020). Post-1950 ages were calibrated with CALIBomb (Reimer et al., 2004) using the Kure Atoll post-bomb marine curve (Andrews et al., 2016).

Four ages from Site 1 and five from Site 2 were incorporated into Bayesian age-depth models using rbacon v2.4.0 in R (Blaauw and Christen, 2011; Blaauw et al., 2020). Default model settings were used except for site-specific accumulation priors selected to match the excavation depths. Baconvergence diagnostics were used to evaluate Markov Chain Monte Carlo mixing. Age estimates were output at 1 cm down-core resolution and interpolated to calculate accumulation rates.

### 2.4. Elemental abundances and sediment analysis

Trace-element abundances were determined for 23 coral-rubble subsamples to evaluate terrigenous inputs, anthropogenic trace metals, environmental variability, and water-composition changes. Samples were processed at the IUI ultraclean laboratory and analysed by ICP-MS at the Institute of Earth Sciences, Hebrew University of Jerusalem, following Chien et al. (2019) and Hartman et al. (2020).

Sediments between coral rubble were collected in 50 mL tubes to minimize loss of fines. Site 1 sediment was collected from 45 and 65 cm excavation depth; Site 2 sediment was collected from 15 and 70 cm. Samples were rinsed with double-distilled water, dried at 80 °C, and wet-sieved through 1000, 500, 250, 125, 100, and 60 µm sieves. The <60 µm fraction was calculated by mass balance. Grain-size statistics were calculated using GRADISTAT (Blott and Pye, 2001), assigning the <60 µm fraction a representative diameter of 10 µm where needed.

### 2.5. Modern coral growth, skeletal density, and calcification

Ten living *A. allingi* colonies were collected near the Site 2 excavation on 18 March 2018, stained for 48 h in seawater containing Alizarin Red-S, and returned unattached to 45 m depth with growth surfaces upward. Colonies were recovered on 16 June 2019, cleaned in 3% sodium hypochlorite, rinsed, dried, and sectioned. Linear extension was measured from the Alizarin band to the new skeletal surface along visually selected corallite growth paths, providing conservative minimum growth estimates (Dustan, 1975).

Skeletal density was measured from 7 mm thick branch slabs using X-radiography with Tridacna-shell density standards and CoralXDS software, following Watanabe et al. (2019). Calcification was calculated as skeletal bulk density multiplied by linear extension, following Lough and Barnes (2000).

### 2.6. Cover-based accretion potential

To evaluate whether present-day coral growth and cover could sustain carbonate accumulation comparable to the radiocarbon-dated rubble record, we estimated a first-order, cover-based potential vertical accretion rate (*A*_mod_, mm yr^−1^) as

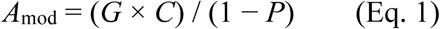

where *G* is the mean linear extension rate of living *A. allingi* colonies (mm yr^−1^; Section 2.5), *C* is fractional live coral cover from the CoralNet photoquadrat surveys (Section 2.1), and *P* is the fractional inter-skeletal space of the rubble deposit (open or sediment-filled voids between skeletal fragments). Assuming that skeleton is added at rate *G* over each colony’s planar footprint (i.e. ignoring colony rugosity and three-dimensional growth form), *G* × *C* is the bulk volume of skeleton added per unit area of seafloor per year, the cover-weighted production term of census-based carbonate budgets (Hubbard et al., 1990), and dividing by (1 − *P*) converts this skeletal volume into deposit thickness. Carbonate-budget studies make this conversion in mass units, dividing carbonate production by aragonite mineral density (≈2.9 g cm^−3^) and by (1 − total porosity) (Smith and Kinsey, 1976; Kinsey, 1985; Kinsey and Hopley, 1991; Dee et al., 2024), which is equivalent only if the total porosity includes the skeleton’s internal porosity. Because *A. allingi* skeletons are highly porous (bulk density 1.17 g cm^−3^, i.e. ≈60% intraskeletal porosity; Section 3.4), applying the 20–50% total porosities commonly assumed for framework reefs (Kinsey, 1985; Browne et al., 2013; Dee et al., 2024) would lower the estimates by a factor of approximately 2.5 for the same porosity value; Eq. (1) is therefore the more generous formulation for testing whether present-day production can account for the observed accumulation. In the absence of direct measurements of inter-skeletal space in rubble beds, we assumed *P* = 0.35 and tested *P* = 0.20–0.50. In situ growth data were obtained only for *A. allingi*, the dominant rubble-forming taxon in the deeper part of both deposits; its extension rate is therefore applied to total live coral cover across all taxa present (including *E. paradivisa*) as a single-species proxy for community-wide linear extension, a simplification discussed further in Section 4.1.

These production-to-accretion conversions, including their porosity assumptions, were developed for shallow-water (mostly <20 m; to 40 m in Hubbard et al., 1990), framework-building coral reefs in the Caribbean and Indo-Pacific/Great Barrier Reef (Smith and Kinsey, 1976; Kinsey, 1985; Hubbard et al., 1990; Browne et al., 2013; Perry et al., 2012, 2013, 2018), not for mesophotic, unconsolidated rubble fields in the Red Sea. Their application here is therefore a first-order, exploratory comparison.

## 3. Results

### 3.1. Modern and fossil coral-rubble assemblages

Modern assemblages differed between sites at 45 m depth. Hermatypic live-coral cover averaged 8.0 ± 3.5% (mean ± 95% confidence interval) at Site 1 and 2.7 ± 1.8% at Site 2, although previous surveys in the region documented local coral cover as high as 30% in some areas (e.g., Eyal-Shaham et al., 2016). The most abundant modern coral categories in the present surveys were the two *Alveopora* spp. and *E. paradivisa* third at Site 1 and *A. allingi* at Site 2. Coral rubble was the dominant lithological component at both sites (Fig. 1).

Excavated rubble showed variable sediment infill and light to moderate alteration through bioerosion, secondary accretion, and cementation, with no simple depth-related preservation trend. Site 1 recorded a pronounced assemblage transition: *E. paradivisa* dominated the upper excavation layers by weight but declined sharply below 35 cm, whereas *A. allingi* dominated deeper layers and live assemblages. At Site 2, *Alveopora* dominated most fossil depth bins to the base of the 70 cm excavation, although *A. ocellata* replaced *A. allingi* in the upper layers (Fig. 2). Minimum rubble thickness was approximately 1 m at Site 1 and 1.5 m at Site 2.

**Figure 2.**
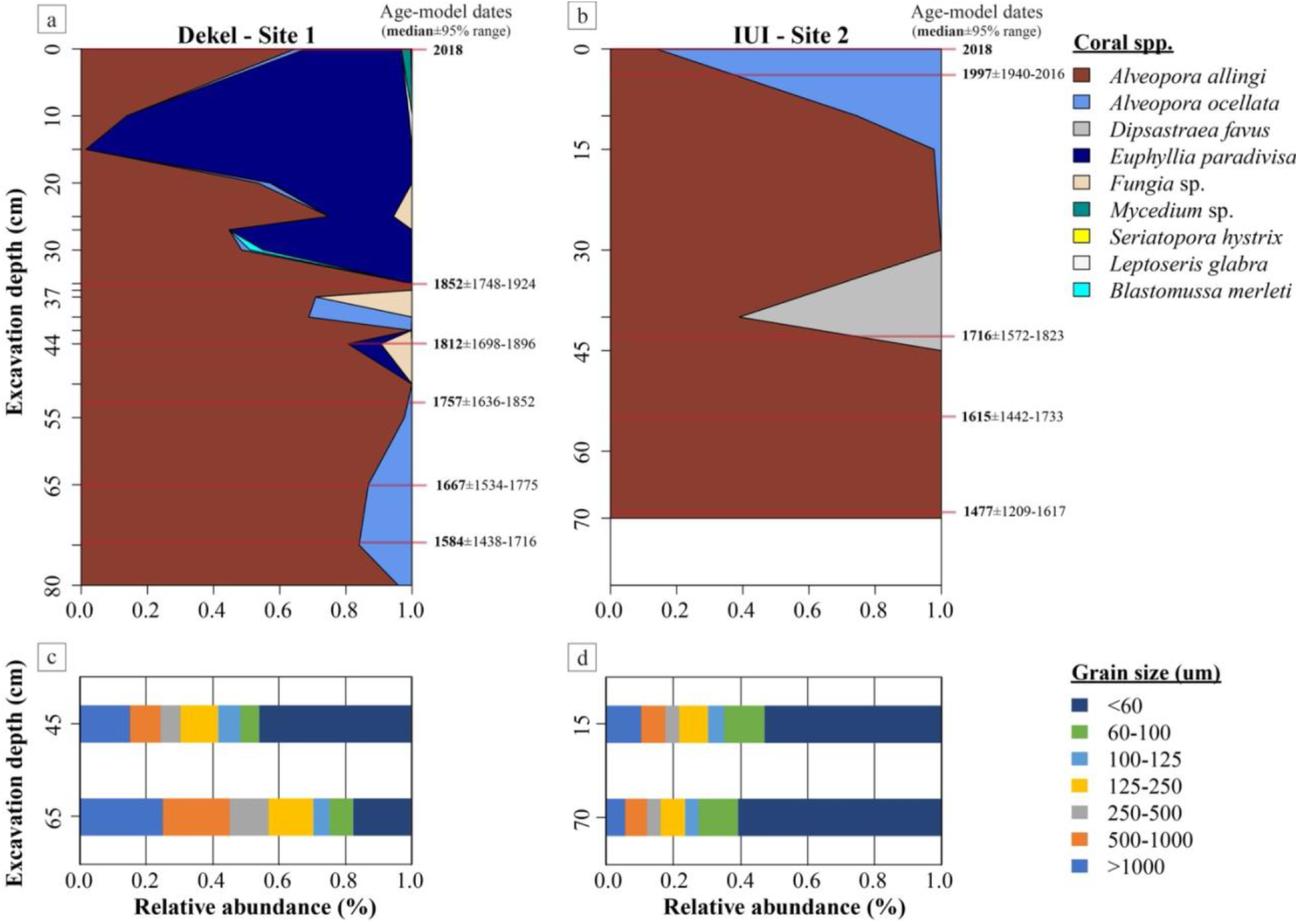
Coral rubble and sediment relative abundances by weight down the excavation pits. (a-b) Colored stacked areas represent coral taxa; red lines indicate depths from which *Alveopora allingi* material was sampled for radiocarbon dating. Age-modelled ranges to the right show AD age ranges and median ages. (c-d) Stacked bars show relative grain-size distributions by weight for sediments collected from two depths within each excavation pit.

### 3.2. Accretion rates

Radiocarbon age-depth models produced near-linear accumulation histories at both sites despite faunal transitions and grain-size variations (Fig. 2; Supplementary Figs. S1, S2). Site 1 accumulated at a mean rate of 2.208 ± 0.520 mm yr^-1^ (mean ± SD), with a 95% modelled range of 1.229-14.286 mm yr^-1^. If the same rate extends to the full measured deposit thickness, Site 1 has accumulated for at least 450 yr. Site 2 accumulated more slowly, with a mean rate of 1.331 ± 0.126 mm yr^-1^ and 95% modelled range of 0.625-6.250 mm yr^-1^. Given a minimum deposit thickness of approximately 1.5 m, Site 2 has likely accumulated for at least 1100 yr.

### 3.3. Sediments and geochemistry

All sediment samples were very poorly sorted. At Site 1, approximately 46% of the 45 cm sample was silt-sized or finer, compared with 18% in the deeper 65 cm sample. The deeper Site 1 sample contained >40% coarse sand or larger grains and had a mean grain size of 346 µm, whereas the shallower Site 1 sample had a mean grain size of 115 µm. Site 2 sediment was finer overall, with 50-60% silt-sized or finer material and mean grain sizes of 82 and 57 µm at 15 and 70 cm, respectively (Fig. 2).

Trace-element data showed enrichment of Pb, V, and Cu in the upper Site 1 layer and also in the deepest Site 1 sample. Site 2 recorded mild enrichments in the upper 20 cm, but when normalized to Al, the uppermost Site 2 sample showed substantial relative enrichment in Pb, V, Cu, Sr, Ba, P, Ti, Nd, Mn, and Mg (Supplementary Figs. S3-S8). These patterns indicate differences in detrital input and upper-section anthropogenic perturbation.

### 3.4. Modern *A. allingi* growth and density

Living *A. allingi* colonies, the major contemporary rubble contributor at both sites, extended at 3.04 ± 0.87 mm yr^-1^, with individual values ranging from 1.59 to 4.97 mm yr^-1^. Skeletal bulk density averaged 1.17 ± 0.27 g cm^-3^ and ranged from 0.75 to 1.67 g cm^-3^. The resulting mean calcification rate was 0.242 ± 0.069 g cm^-2^ yr^-1^ (Fig. 3), calcification was computed per colony (n = 4) for the subset with both measurements.

**Figure 3.**
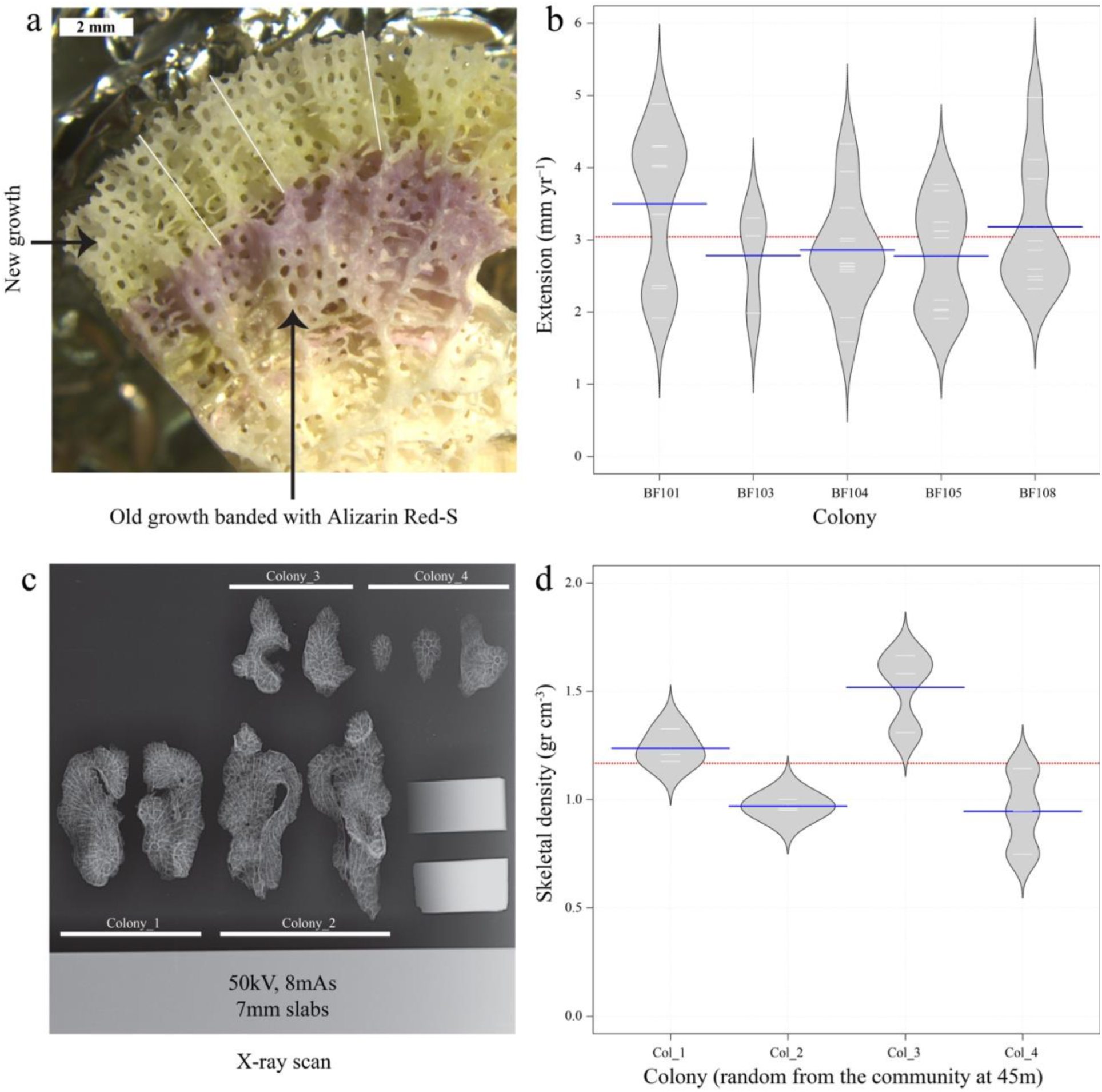
In situ growth and skeletal density of modern *Alveopora allingi* colonies, the major contemporary contributor to rubble deposits at 45 m depth in Eilat. (a) Skeleton slab cross-section with Alizarin Red-S band and new growth; white crosslines indicate linear-extension measurements. (b,d) Bean plots showing means as blue crosslines, individual data as white lines, estimated density as polygons, and overall mean as red dashed line. (c) X-ray scan of 7 mm skeletal slabs from four colonies with density standards.

## 4. Discussion

### 4.1. Mesophotic rubble fields as active carbonate factories

The central finding of this study is that Red Sea mesophotic coral-rubble fields can sustain positive late Holocene carbonate accumulation. The mean rates of 1.33-2.21 mm yr^-1^ are higher than the approximately 0.3 mm yr^-1^ measured from short mesophotic Caribbean cores by Sherman et al. (2023) and overlap or exceed some modern carbonate-budget estimates from deeper shallow-water reefs (Perry et al., 2013). The deposits also exceed 1-1.5 m in thickness, demonstrating that these habitats are not merely thin surficial veneers over older hardgrounds. Regional comparisons emphasize that the result is important because of its depositional style and depth rather than because it exceeds shallow reef-crest production. Published Red Sea Holocene accretion rates include values around 0.67–1.5 mm yr^−1^ from earlier reef studies (Dullo et al., 1996; Purkis et al., 2010) and 3.6–9.3 mm yr^−1^ from recent lagoonal and fore-reef cores in Eilat (Feldman et al., 2025). In shallow (5–8 m) reef-slope rubble from the central and southern Red Sea, Hammerman et al. (2022) reported 18 core-averaged accumulation rates ranging from 0.45 to 71.6 mm yr^−1^ (median 8.1 mm yr^−1^; mean 12.5 mm yr^−1^, or 9.0 mm yr^−1^ excluding the extreme value from core AS4). The present mesophotic rubble rates sit near the lower end of that regional distribution but overlap low-accretion Red Sea cores (1.44-1.89 mm yr^-1^). Thus, the GoEA mesophotic coral-rubble fields (CRFs) are slower than most shallow/core framework examples, but they are nevertheless persistent, metre-scale, actively accumulating carbonate bodies at 40-65 m depth.

The modern growth and cover data suggest that present living coral cover alone cannot account for the radiocarbon-derived accumulation. Applying Eq. (1) with the measured *A. allingi* extension rate (3.04 mm yr^−1^), present cover and *P* = 0.35 yields potential accretion of only approximately 0.37 mm yr^−1^ at Site 1 and 0.13 mm yr^−1^ at Site 2 (0.30–0.49 and 0.10–0.16 mm yr^−1^ for *P* = 0.20–0.50), approximately 6-fold (4.5–7.3) and 10-fold (8–13) below the respective long-term age-depth rates; closing this gap at present cover would require implausibly high inter-skeletal space (*P* ≈ 0.89 and 0.94). At the approximately 30% cover reported previously from the region (Section 3.1), the estimate (1.40 mm yr^−1^; 1.14–1.82 mm yr^−1^ for *P* = 0.20–0.50) approaches or exceeds the long-term rate of Site 2 but remains approximately 1.6-fold (1.2–1.9) below that of Site 1. Matching the long-term rates through in situ coral production alone would require approximately 47% cover at Site 1 and 28% at Site 2 (*P* = 0.35), roughly 6- and 10-times present cover and, at Site 1, more than the ≈33–34% total cover reported at 40–60 m by Eyal-Shaham et al. (2016). Because Eq. (1) assumes complete retention of skeletal production, closing the gap requires processes that add volume, such as historically higher coral cover, carbonate from other calcifiers and secondary calcification, and sediment trapped beyond the inter-skeletal space; mortality, post-mortem loss and reworking would only widen it. The comparison therefore supports a process interpretation in which CRF accretion is not a direct snapshot of live coral cover but a time-integrated balance among skeletal production, retention, porosity, cementation, and loss. More broadly, coral linear extension rates can be decoupled from reef vertical accretion rates over Holocene timescales (Dullo, 2005; Roff, 2020). Woodroffe and Webster (2014) explicitly distinguish individual coral linear extension (up to 100 mm yr^−1^ in branching corals and 10–20 mm yr^−1^ in large domal corals) from whole-reef vertical accretion, which integrates porosity, framework construction, and community composition. Here, this decoupling is compounded by using the *A. allingi* extension rate as a proxy for all cover-forming taxa (Section 2.6), as it is the only taxon with in situ growth data; other rubble-forming corals (e.g., *E. paradivisa*) may differ in extension rate and growth form, so this proxy could bias the cover-based estimate in either direction. Notably, the mismatch runs opposite to the usual timescale dependence of accumulation rates, whereby short-term rates exceed long-term averages because the latter integrate hiatuses and erosion (Sadler, 1981); the shortfall of the cover-based estimate is therefore unlikely to be an artefact of the different averaging intervals.

Unlike shallow reefs, whose accretion potential is usually assessed against projected sea-level rise (Perry et al., 2018), rubble fields at 40–65 m depth are not limited by accommodation space; the comparison above therefore tests whether present-day production is consistent with the deposits’ own long-term accumulation rather than their capacity to track sea level. Because the underlying production-to-accretion framework was developed for shallow-water reefs (Section 2.6), this comparison is first-order; direct measurements of rubble-bed porosity, cementation and burial history (Section 4.4) are needed to make it more quantitative.

### 4.2. Taphonomy and depositional continuity

Taphonomic bias is a critical uncertainty when interpreting rubble assemblages. Branching or phaceloid corals can be overrepresented in death assemblages because they fragment readily, and rubble can be transported or mixed during storms, gravity-driven downslope movement, or bioturbation (Pandolfi and Greenstein, 1997; Rasser and Riegl, 2002). In this study, however, several observations support a dominantly local, active depositional interpretation. First, the modern living assemblages contain the same broad rubble-forming taxa found in the excavations. Second, abundant dead rubble occurs adjacent to living colonies, consistent with continuous in situ skeletal turnover. Third, age-depth models show broadly coherent accumulation histories, and minor age reversals are consistent with localized mixing rather than catastrophic remobilization of the entire deposit.

The rubble fields therefore represent a distinct mode of reef accretion. They lack the rigid, high-relief framework commonly associated with shallow-water reef growth, but they retain carbonate through interlocking skeletal fragments, trapped sediment, secondary calcification, and partial cementation. Such low-relief depositional systems can build substantial thickness if skeletal residence time and retention exceed physical and biological loss. This mechanism provides a modern analogue for interpreting poorly lithified coral-rubble deposits in both Holocene and older records.

### 4.3. Faunal transitions and environmental signals

The rubble assemblages record ecological transitions that are not evident from the living surface alone. At Site 1, the sharp decline of *E. paradivisa* below 35 cm and dominance of *A. allingi* at depth suggest a local community shift. At Site 2, persistence of *Alveopora* through the excavated interval indicates greater faunal continuity, although upper layers record replacement of *A. allingi* by *A. ocellata*. These differences demonstrate that adjacent mesophotic CRFs can record different ecological trajectories over centennial to millennial timescales.

Sediment and elemental data provide additional environmental context. Finer sediment at Site 2 is consistent with the flat terrace setting and reduced hydrodynamic winnowing relative to Site 1, whereas the upward fining at Site 1 may reflect changes in terrigenous delivery, local hydrodynamics, or rubble baffling as the deposit developed. Enrichment of Pb, V, Cu, and other elements in upper sections is consistent with recent anthropogenic inputs superimposed on natural detrital variability, especially when evaluated relative to Al. The deepest Site 1 enrichment suggests that similar elemental patterns can also arise from older non-anthropogenic or diagenetic processes, reinforcing the need to interpret trace elements with age control and detrital normalization.

### 4.4. Implications for mesophotic reef development

These results revise how mesophotic reef geomorphology should be conceptualized in the Red Sea and similar settings. MCEs can be active carbonate factories even when they form unconsolidated, low-relief rubble fields rather than cemented reef framework. Their deposits record long-term ecological baselines, sedimentary regime shifts, and geochemical perturbations across depths that are rarely sampled by conventional reef coring. Because shallow reef palaeoecological records dominate current historical baselines, mesophotic rubble deposits can extend reconstructions into deeper, light-dependent parts of reef systems.

The findings also have conservation relevance as shallow reefs face intensifying climate and anthropogenic stress (Hughes et al., 2003; Reimer et al., 2024). Mesophotic CRFs are often treated as marginal habitats, but the observed accretion, live coral growth, and palaeoecological archive show that they contribute to regional carbonate budgets and habitat persistence. Future work should quantify spatial extent, measure porosity and secondary cementation directly, expand multi-species calcification budgets, and test whether similar accretionary rubble systems occur in other Red Sea and Indo-Pacific mesophotic settings.

## 5. Conclusions

Diver excavations and radiocarbon age-depth models show that northern Red Sea mesophotic CRFs are metre-scale late Holocene carbonate deposits rather than thin biological veneers. Deposits accumulated at mean rates of 1.33-2.21 mm yr^-1^ and record faunal, sedimentological, and geochemical change over centennial to millennial timescales. Modern *A. allingi* growth is substantial for mesophotic depth, but first-order cover-based accretion estimates at present cover underpredict the long-term rates, indicating that historical cover, secondary accretion and sediment retention are essential to rubble-field development. Mesophotic CRFs should therefore be recognized as active carbonate factories and palaeoecological archives within reef-margin systems.

## CRediT authorship contribution statement

Conceptualization: G.E., D.K.W.; Methodology: G.E., D.K.W., N.M.H., Y.S.; Investigation: G.E., D.K.W., Y.S., A.T.; Visualization: G.E., D.K.W., N.M.H.; Funding acquisition: D.K.W., A.T.; Supervision: G.E., A.T.; Writing - original draft: G.E., D.K.W.; Writing - review and editing: all authors.

## Data availability

All codes, radiocarbon, geochemical, sediment, and growth datasets will be made available in an appropriate public repository upon acceptance.

## Declaration of generative AI and AI-assisted technologies in the writing process

During the preparation of this work, the authors used Claude (Anthropic) for journal-specific formatting and language editing. After using these tools, the authors reviewed and edited the content as needed, verified all references against the original sources, and take full responsibility for the content of the published article.

## Acknowledgments

We thank the Interuniversity Institute for Marine Sciences in Eilat, associated students and staff, E. Sestieri for boat support, R. Tamir for support with closed-circuit rebreather diving, accretion-thickness surveys, drop-camera surveys, coral-cover surveys, and consultation, and O. Hameiri, J. Bellworthy, and G. Banc-Prandi for open-circuit diving support.

