## Supplementary materials for "Active late Holocene accretion of mesophotic coral-rubble fields in the northern Red Sea"

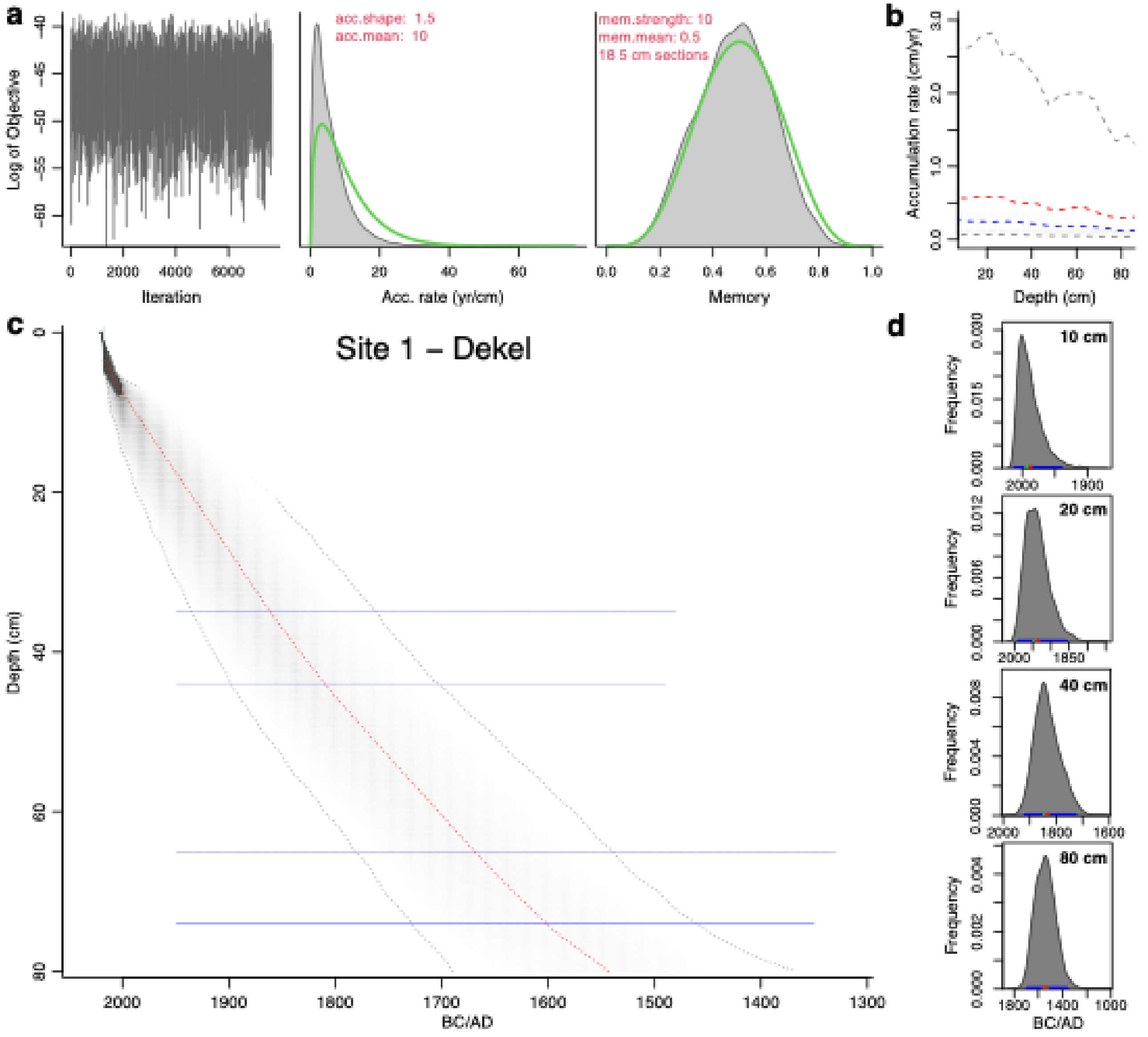


*Figure S1:* Bayesian age-depth modeling results for Site 1 – Dekel's pit.
*(a) MCMC optimization diagnostics showing log objective values across iterations, indicating convergence. Posterior distribution of accumulation rate (cm yr$^{-1}$) with acc.shape = 1.5 and acc.mean = 10. Posterior distribution of memory parameters with mem.strength = 10 and mem.mean = 0.5.*
*(b) Posterior distribution of accumulation rate profile (cm yr$^{-1}$) with dashed red and blue lines indicating the mean and median accumulation rates, respectively.* **.** *Gray dashed lines represent the 95% confidence interval, capturing the uncertainty range around the estimated accumulation rates at each depth.*
*(c) Age-depth model with median age estimates (black line) and 95% confidence intervals (gray envelope) across 18 sections of 5 cm.*
*(d) Posterior age distributions for selected pit depths (10 cm, 20 cm, 40 cm, and 80 cm) showing age uncertainty at each depth.*

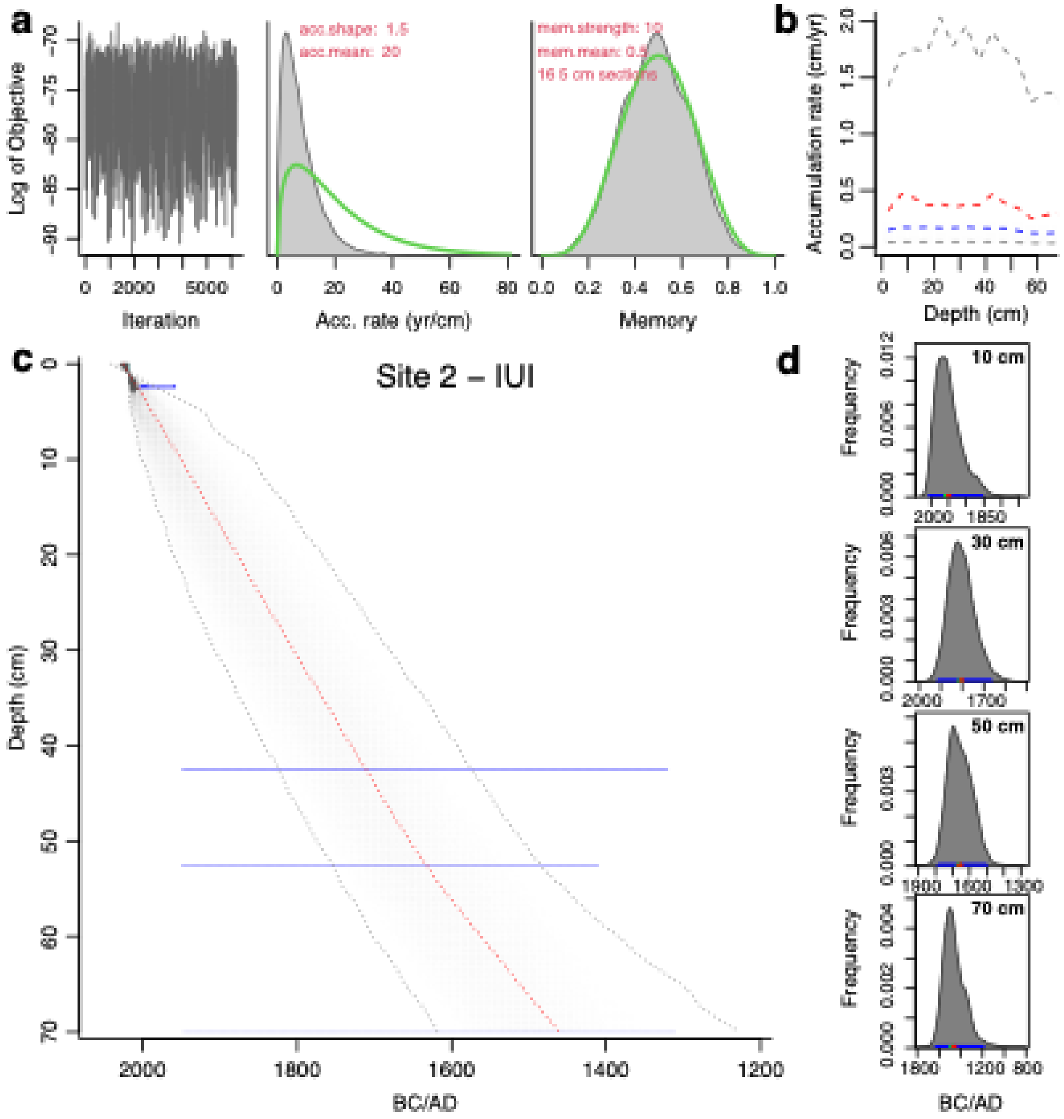


*Figure S2:* Bayesian age-depth modeling results for Site 2 – IUI Site 2 pit.
*(a) MCMC optimization diagnostics showing log objective values across iterations, indicating convergence. Posterior distribution of accumulation rate (cm yr$^{-1}$) with acc.shape = 1.5 and acc.mean = 20. Posterior distribution of memory parameters with mem.strength = 10 and mem.mean = 0.5.*
*(b) Posterior distribution of accumulation rate profile (cm yr$^{-1}$) with dashed red and blue lines indicating the mean and median accumulation rates, respectively.* **.** *Gray dashed lines represent the 95% confidence interval, capturing the uncertainty range around the estimated accumulation rates at each depth.*
*(c) Age-depth model with median age estimates (black line) and 95% confidence intervals (gray envelope) across 16 sections of 5 cm.*
*(d) Posterior age distributions for selected pit depths (10 cm, 30 cm, 50 cm, and 70 cm) showing age uncertainty at each depth.*

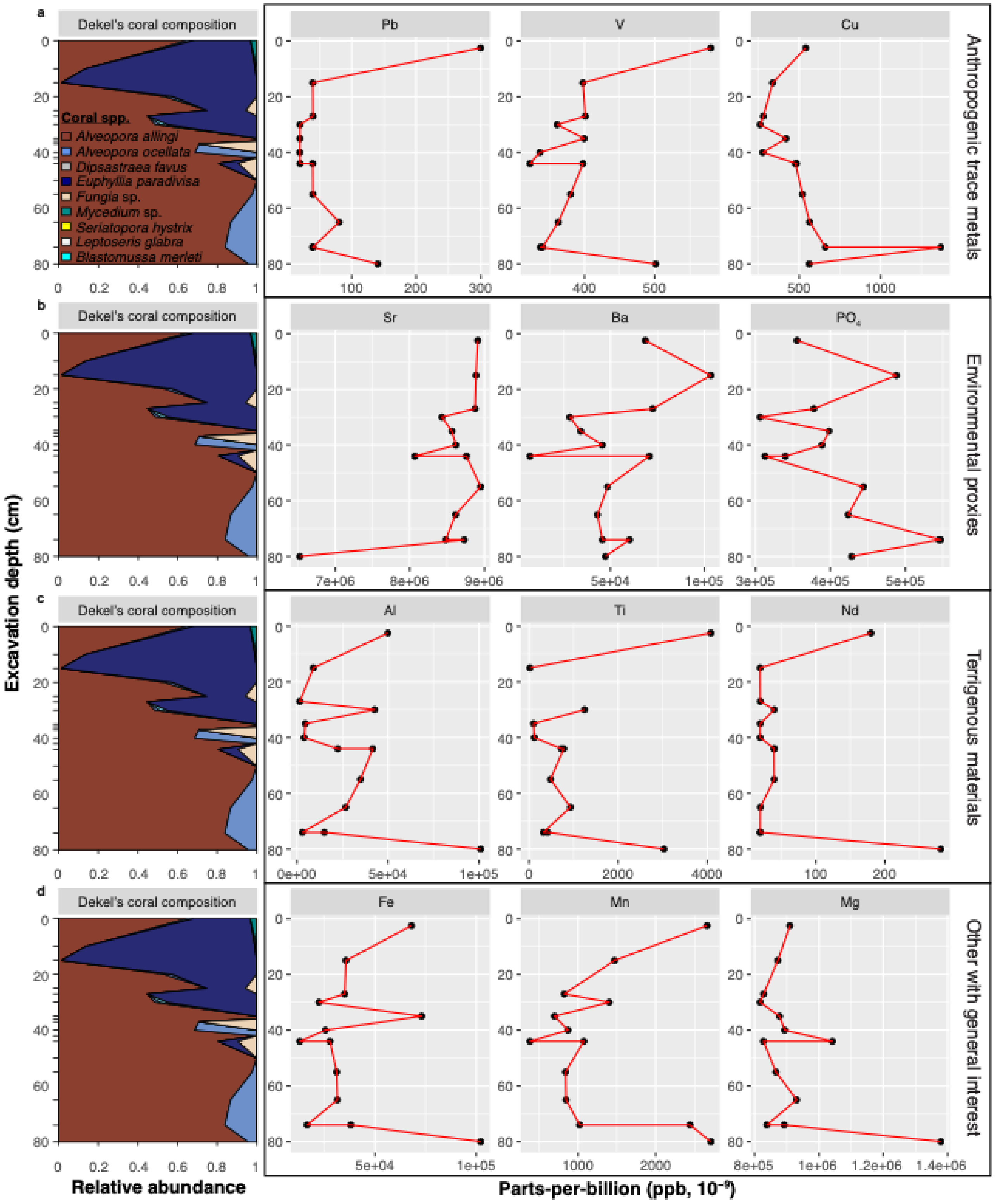


*Figure S3: Site 1 – Non-transformed geochemical composition. Geochemical and ecological stratigraphy of Dekel Site 1 pit samples. Relative abundance of fossil coral taxa across pit depths, showing species composition changes over time. (a-d) Concentrations of anthropogenic trace metals (e.g., Pb, V, Cu), environmental proxies (e.g., Sr, Ba, $PO_4$), and terrigenous materials (e.g., Al, Ti, Nd) along excavation depth (cm), expressed in parts-per-billion (ppb).*

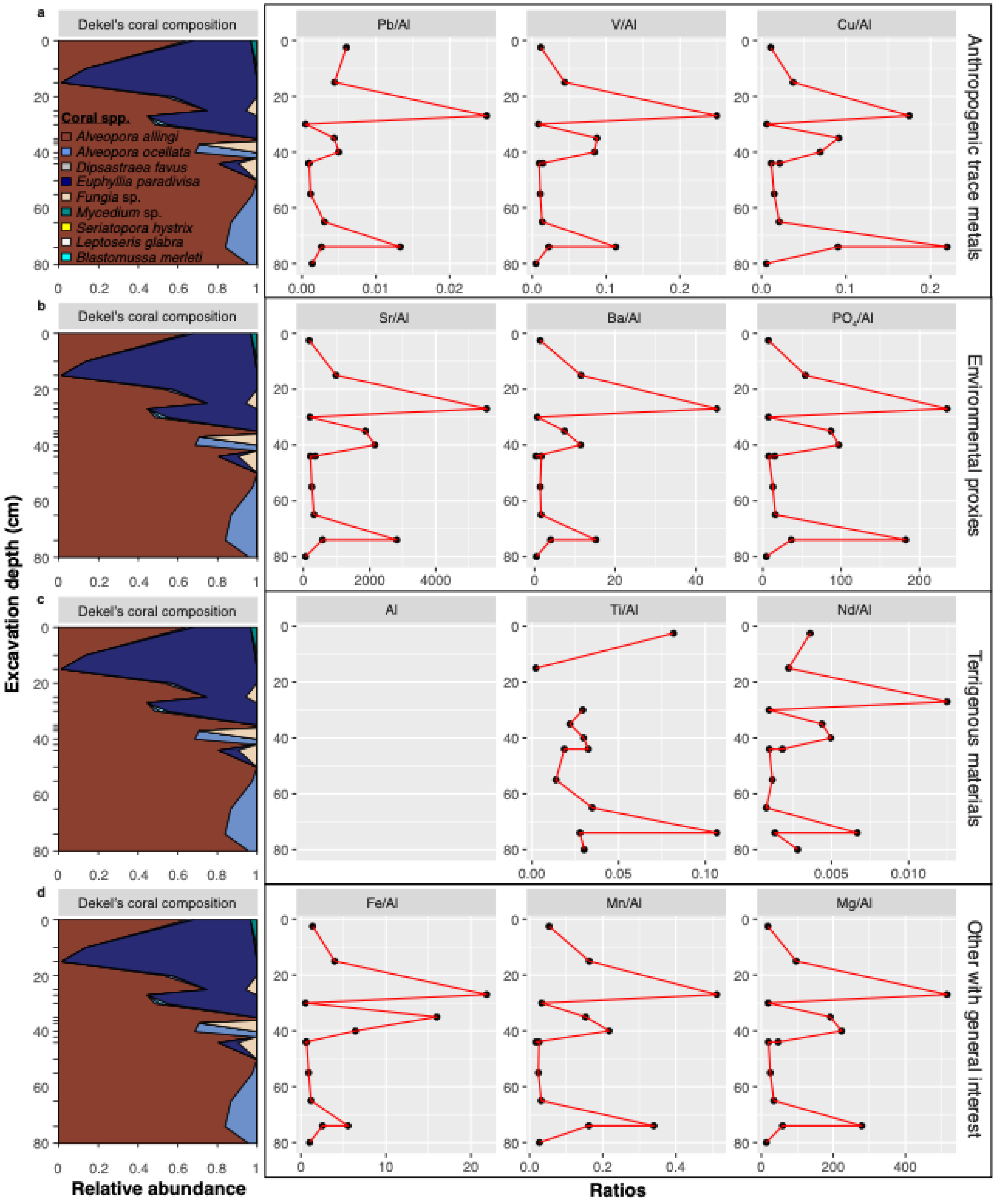


*Figure S4: Site 1 – Al-transformed geochemical composition. Geochemical and ecological stratigraphy of Dekel Site 1 pit samples. Relative abundance of fossil coral taxa across pit depths, showing species composition changes over time. (a-d) Concentrations of anthropogenic trace metals (e.g., Pb, V, Cu), environmental proxies (e.g., Sr, Ba, $PO_4$), and terrigenous materials (e.g., Al, Ti, Nd) along excavation depth (cm), expressed in parts-per-billion (ppb).*

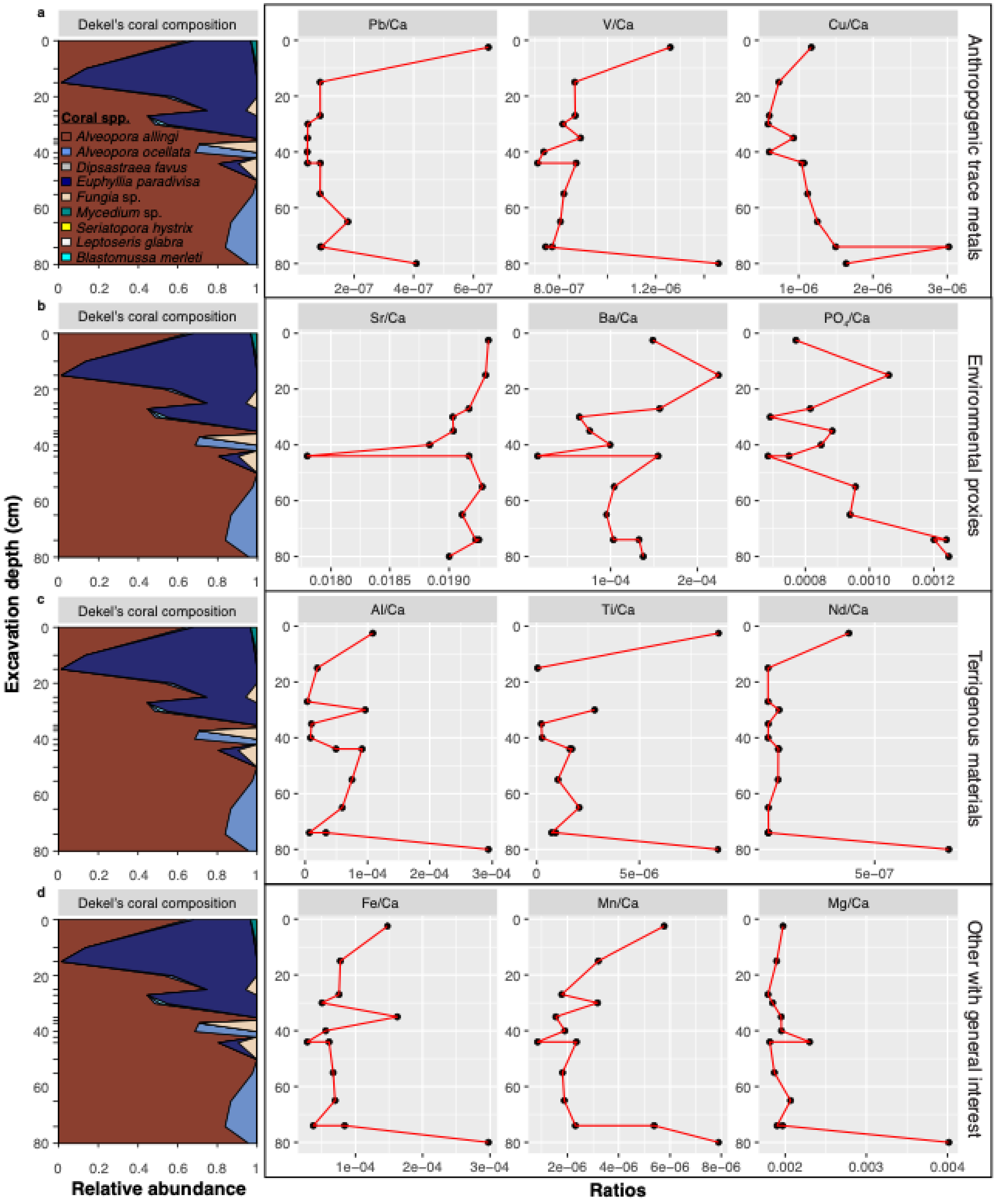


*Figure S5: Site 1 – $^{44}$Ca-transformed geochemical composition. Geochemical and ecological stratigraphy of Dekel Site 1 pit samples. Relative abundance of fossil coral taxa across pit depths, showing species composition changes over time. (a-d) Concentrations of anthropogenic trace metals (e.g., Pb, V, Cu), environmental proxies (e.g., Sr, Ba, $PO_4$), and terrigenous materials (e.g., Al, Ti, Nd) along excavation depth (cm), expressed in parts-per-billion (ppb).*

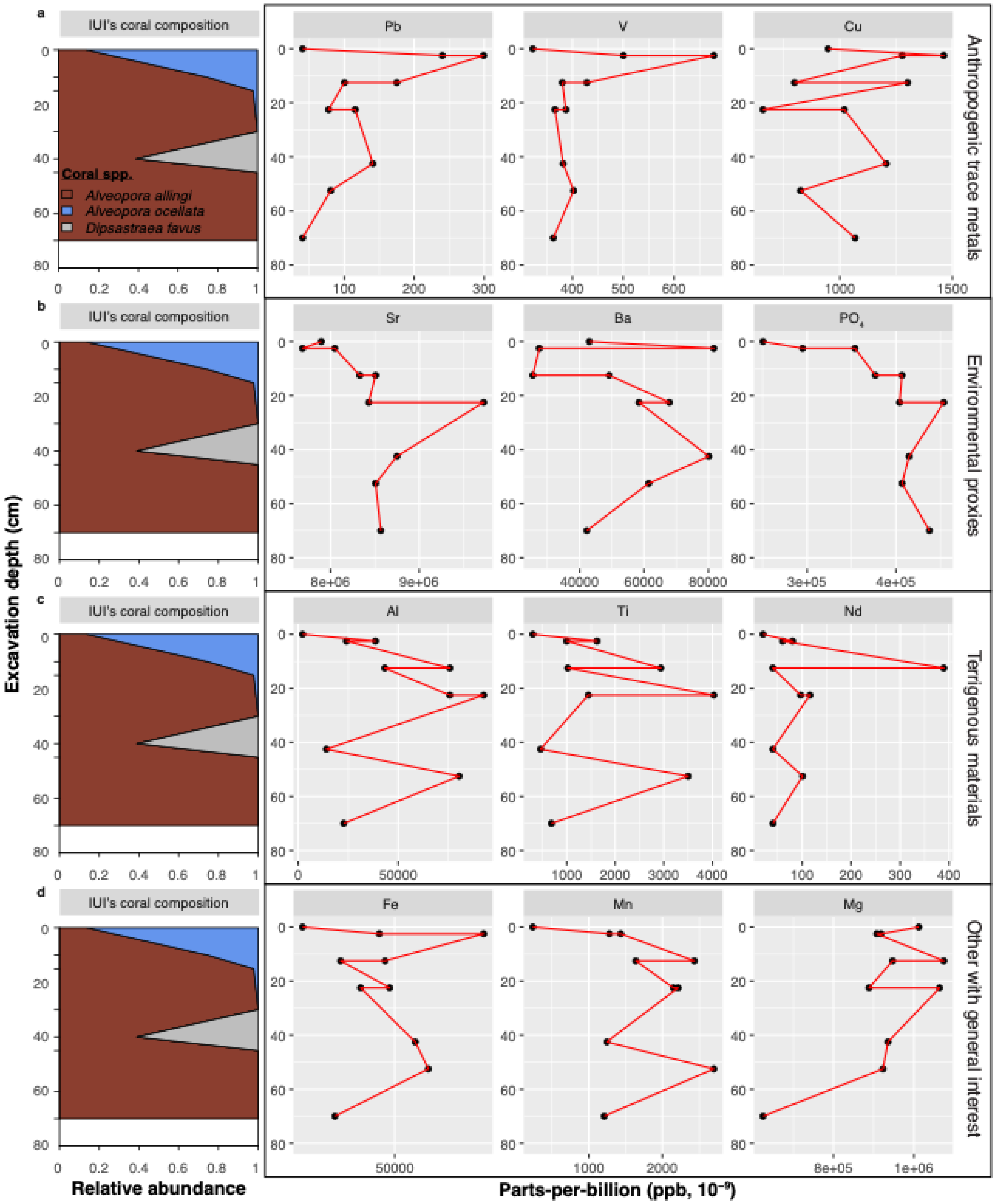


*Figure S6: Site 2 – Non-transformed geochemical composition. Geochemical and ecological stratigraphy of IUI Site 2 pit samples. Relative abundance of fossil coral taxa across pit depths, showing species composition changes over time. (a-d) Concentrations of anthropogenic trace metals (e.g., Pb, V, Cu), environmental proxies (e.g., Sr, Ba, $PO_4$), and terrigenous materials (e.g., Al, Ti, Nd) along excavation depth (cm), expressed in parts-per-billion (ppb).*

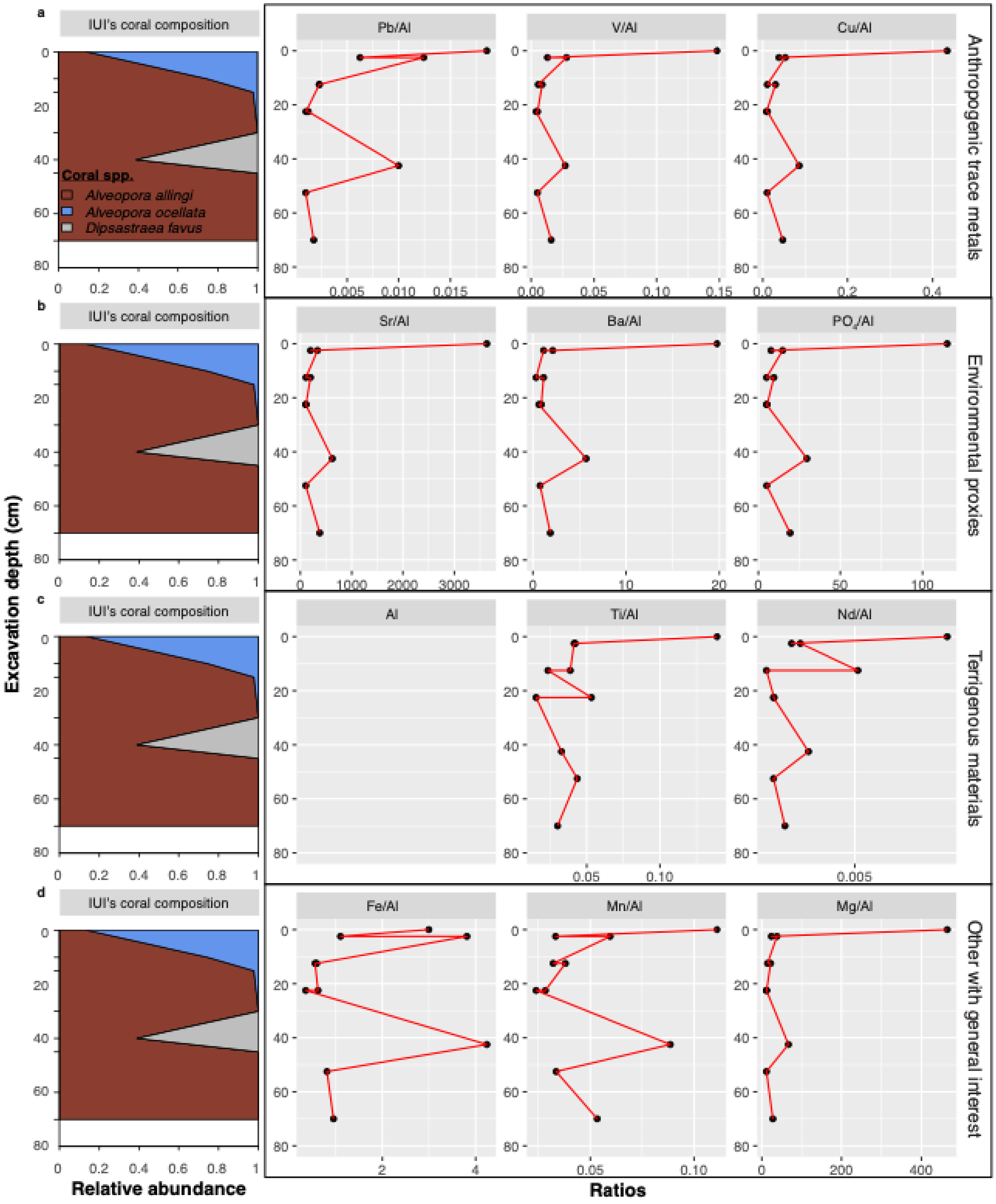


*Figure S7: Site 2 – Al-transformed geochemical composition. Geochemical and ecological stratigraphy of IUI Site 2 pit samples. Relative abundance of fossil coral taxa across pit depths, showing species composition changes over time. (a-d) Concentrations of anthropogenic trace metals (e.g., Pb, V, Cu), environmental proxies (e.g., Sr, Ba, $PO_4$), and terrigenous materials (e.g., Al, Ti, Nd) along excavation depth (cm), expressed in parts-per-billion (ppb).*

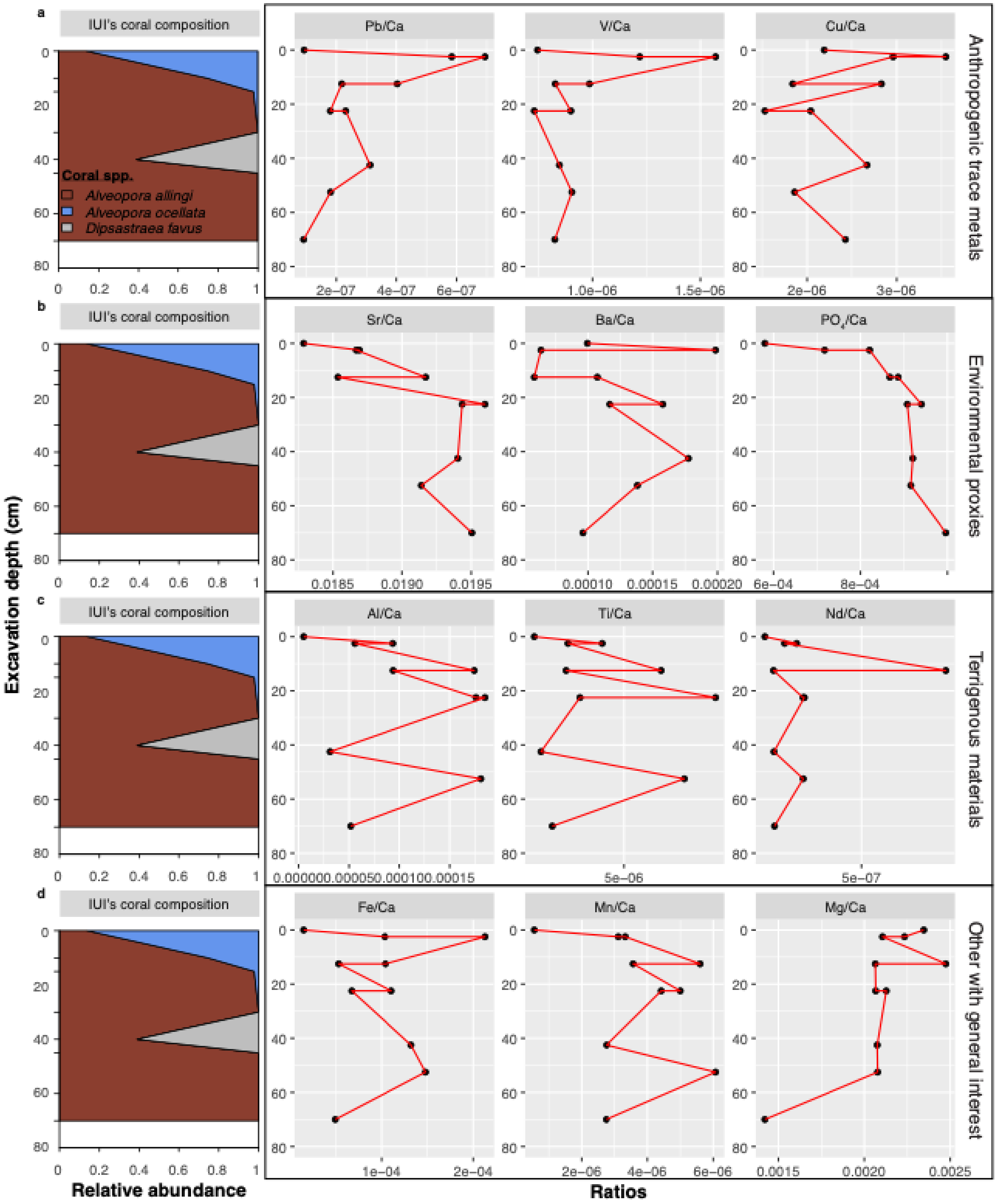


*Figure S8: Site 2 – $^{44}$Ca-transformed geochemical composition. Geochemical and ecological stratigraphy of IUI Site 2 pit samples. Relative abundance of fossil coral taxa across pit depths, showing species composition changes over time. (a-d) Concentrations of anthropogenic trace metals (e.g., Pb, V, Cu), environmental proxies (e.g., Sr, Ba, $PO_4$), and terrigenous materials (e.g., Al, Ti, Nd) along excavation depth (cm), expressed in parts-per-billion (ppb).*

Table S1: Granular sediment grain-size statistics

| | | Dekel_45cm | Dekel_65cm | IUI_15cm | IUI_70cm |
|---|---|---|---|---|---|
| | ANALYST AND DATE: | G, Eyal, | G, Eyal, | G, Eyal, | G, Eyal, |
| | SIEVING ERROR: | | | | |
| | SAMPLE TYPE: | Trimodal, Very Poorly Sorted | Bimodal, Very Poorly Sorted | Bimodal, Very Poorly Sorted | Bimodal, Very Poorly Sorted |
| | TEXTURAL GROUP: | Gravelly Mud | Gravelly Muddy Sand | Gravelly Mud | Slightly Gravelly Sandy Mud |
| | SEDIMENT NAME: | Fine Gravelly Coarse Silt | Fine Gravelly Very Coarse Silty Coarse Sand | Fine Gravelly Coarse Silt | Slightly Fine Gravelly Very Fine Sandy Coarse Silt |
| METHOD OF MOMENTS Arithmetic (∝m) | MEAN ($\bar{x}_a$): | 821.4 | 1366.0 | 592.1 | 370.5 |
| | SORTING ($\sigma_a$): | 1565.1 | 1829.2 | 1353.4 | 1034.9 |
| | SKEWNESS ($Sk_a$): | 1.881 | 1.090 | 2.486 | 3.625 |
| | KURTOSIS ($K_a$): | 4.647 | 2.273 | 7.337 | 14.56 |
| METHOD OF MOMENTS Geometric (∝m) | MEAN ($\bar{x}_g$): | 119.2 | 331.5 | 84.00 | 62.61 |
| | SORTING ($\sigma_g$): | 5.749 | 5.201 | 4.955 | 4.062 |
| | SKEWNESS ($Sk_g$): | 0.670 | -0.190 | 1.083 | 1.412 |
| | KURTOSIS ($K_g$): | 2.040 | 1.871 | 2.893 | 3.948 |
| METHOD OF MOMENTS Logarithmic (ϕ) | MEAN ($\bar{x}_\phi$): | 3.069 | 1.593 | 3.574 | 3.997 |
| | SORTING ($\sigma_\phi$): | 2.523 | 2.379 | 2.309 | 2.022 |
| | SKEWNESS ($Sk_\phi$): | -0.670 | 0.190 | -1.083 | -1.412 |
| | KURTOSIS ($K_\phi$): | 2.040 | 1.871 | 2.893 | 3.948 |
| FOLK AND WARD METHOD (∝m) | MEAN ($M_G$): | 115.3 | 345.6 | 81.97 | 57.20 |
| | SORTING ($\sigma_G$): | 6.407 | 6.061 | 5.570 | 4.130 |
| | SKEWNESS ($Sk_G$): | 0.264 | -0.078 | 0.394 | 0.361 |
| | KURTOSIS ($K_G$): | 0.828 | 1.029 | 1.072 | 1.138 |
| FOLK AND WARD METHOD (ϕ) | MEAN ($M_Z$): | 3.117 | 1.533 | 3.609 | 4.128 |
| | SORTING ($\sigma_I$): | 2.680 | 2.600 | 2.478 | 2.046 |
| | SKEWNESS ($Sk_I$): | -0.264 | 0.078 | -0.394 | -0.361 |
| | KURTOSIS ($K_G$): | 0.828 | 1.029 | 1.072 | 1.138 |
| FOLK AND WARD METHOD (Description) | MEAN: | Very Fine Sand | Medium Sand | Very Fine Sand | Very Coarse Silt |
| | SORTING: | Very Poorly Sorted | Very Poorly Sorted | Very Poorly Sorted | Very Poorly Sorted |
| | SKEWNESS: | Coarse Skewed | Symmetrical | Very Coarse Skewed | Very Coarse Skewed |
| | KURTOSIS: | Platykurtic | Mesokurtic | Mesokurtic | Leptokurtic |
| | MODE 1 (∝m): | 112.5 | 750.0 | 35.00 | 35.00 |
| | MODE 2 (∝m): | 35.00 | 112.5 | 750.0 | 750.0 |
| | MODE 3 (∝m): | 750.0 | | | |
| | MODE 1 (ϕ): | 3.161 | 0.500 | 5.351 | 5.351 |
| | MODE 2 (ϕ): | 5.351 | 3.161 | 0.500 | 0.500 |
| | MODE 3 (ϕ): | 0.500 | | | |
| | $D_{10}$ (∝m): | 14.77 | 27.85 | 14.06 | 13.44 |
| | $D_{50}$ (∝m): | 87.76 | 377.9 | 54.86 | 43.80 |
| | $D_{90}$ (∝m): | 2018.2 | 3489.7 | 1109.6 | 628.1 |
| | ($D_{90}$ / $D_{10}$) (∝m): | 136.6 | 125.3 | 78.94 | 46.74 |
| | ($D_{90}$ - $D_{10}$) (∝m): | 2003.4 | 3461.9 | 1095.5 | 614.6 |
| | ($D_{75}$ / $D_{25}$) (∝m): | 17.63 | 9.897 | 8.268 | 5.486 |
| | ($D_{75}$ - $D_{25}$) (∝m): | 441.1 | 903.8 | 170.2 | 93.89 |
| | $D_{10}$ (ϕ): | -1.013 | -1.803 | -0.150 | 0.671 |
| | $D_{50}$ (ϕ): | 3.510 | 1.404 | 4.188 | 4.513 |
| | $D_{90}$ (ϕ): | 6.081 | 5.166 | 6.153 | 6.218 |
| | ($D_{90}$ / $D_{10}$) (ϕ): | -6.003 | -2.865 | -41.018 | 9.267 |
| | ($D_{90}$ - $D_{10}$) (ϕ): | 7.094 | 6.969 | 6.303 | 5.547 |
| | ($D_{75}$ / $D_{25}$) (ϕ): | 4.775 | -423.614 | 2.287 | 1.786 |
| | ($D_{75}$ - $D_{25}$) (ϕ): | 4.140 | 3.307 | 3.048 | 2.456 |
| | % GRAVEL: | 10.1% | 16.7% | 7.0% | 3.8% |
| | % SAND: | 43.6% | 65.2% | 39.4% | 34.6% |
| | % MUD: | 46.4% | 18.1% | 53.6% | 61.6% |
| | % V COARSE GRAVEL: | 0.0% | 0.0% | 0.0% | 0.0% |
| | % COARSE GRAVEL: | 0.0% | 0.0% | 0.0% | 0.0% |
| | % MEDIUM GRAVEL: | 0.0% | 0.0% | 0.0% | 0.0% |
| | % FINE GRAVEL: | 5.0% | 8.4% | 3.5% | 1.9% |
| | % V FINE GRAVEL: | 5.0% | 8.4% | 3.5% | 1.9% |
| | % V COARSE SAND: | 5.0% | 8.4% | 3.5% | 1.9% |
| | % COARSE SAND: | 9.3% | 20.2% | 7.0% | 6.4% |
| | % MEDIUM SAND: | 6.1% | 11.7% | 4.4% | 4.3% |
| | % FINE SAND: | 11.2% | 13.4% | 8.3% | 7.0% |
| | % V FINE SAND: | 11.9% | 11.6% | 16.1% | 15.1% |
| | % V COARSE SILT: | 17.2% | 6.9% | 20.1% | 23.0% |
| | % COARSE SILT: | 17.8% | 6.8% | 20.4% | 23.5% |
| | % MEDIUM SILT: | 11.4% | 4.4% | 13.1% | 15.1% |
| | % FINE SILT: | 0.0% | 0.0% | 0.0% | 0.0% |
| | % V FINE SILT: | 0.0% | 0.0% | 0.0% | 0.0% |
| | % CLAY: | 0.0% | 0.0% | 0.0% | 0.0% |

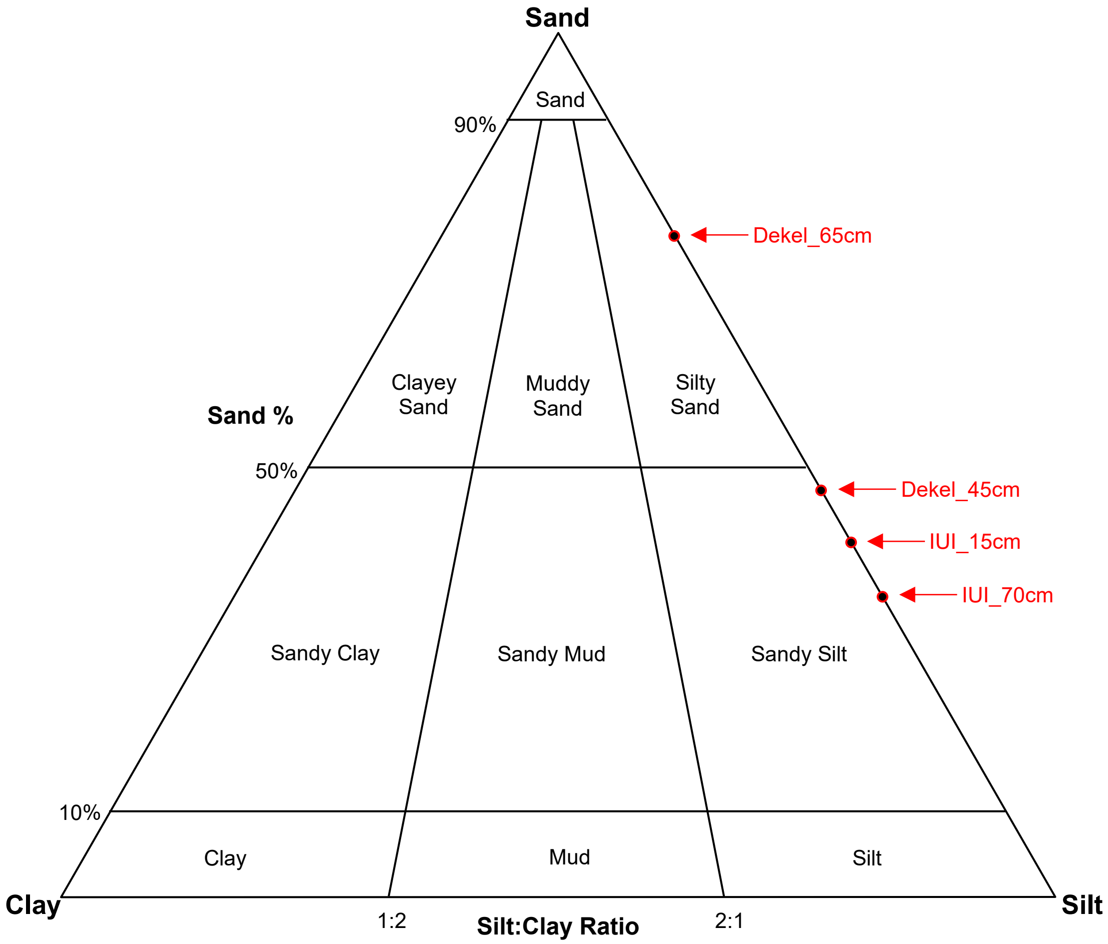


*Figure S9: Distribution of grain-size in different pit's depth plotted on the Sand-Silt-Clay ternary diagram.*
